# Dismantling and rebuilding the trisulfide cofactor demonstrates its essential role in human sulfide quinone oxidoreductase

**DOI:** 10.1101/2020.05.19.103010

**Authors:** Aaron P. Landry, Sojin Moon, Jenner Bonanata, Uhn Soo Cho, E. Laura Coitiño, Ruma Banerjee

## Abstract

Sulfide quinone oxidoreductase (SQR) catalyzes the first step in sulfide clearance, coupling H_2_S oxidation to coenzyme Q reduction. Recent structures of human SQR revealed a sulfur atom bridging the SQR active site cysteines in a trisulfide configuration. Here, we assessed the importance of this cofactor using kinetic, crystallographic and computational modeling approaches. Cyanolysis of SQR proceeds via formation of an intense charge transfer complex that subsequently decays to eliminate thiocyanate. Cyanolysis leads to reversible loss of SQR activity, which is restored in the presence of sulfide. We captured a crystallographic intermediate in SQR that provides clues as to how the oxidized state of the cysteines is preserved. Computational modeling and MD simulations revealed an ~10^5^-fold rate enhancement for nucleophilic addition of sulfide into the trisulfide versus a disulfide cofactor. The cysteine trisulfide in SQR is thus critical for activity and provides a significant catalytic advantage over a cysteine disulfide.

## Introduction

Hydrogen sulfide (H_2_S)^1^ is a signaling molecule that exerts physiological effects in the cardiovascular, central nervous, and gastrointestinal systems (1–3). H_2_S is synthesized endogenously in mammals through the activities of cystathionine β-synthase (4) and cystathionine γ-lyase (5), as well as 3-mercaptopyruvate sulfur transferase (6,7). Tissue concentrations of H_2_S typically range from 10-80 nM (8–10). At higher concentrations, H_2_S can act as a respiratory poison that blocks the electron transport chain by inhibiting complex IV (11).

Due to the bimodal effects of H_2_S, its levels must be strictly regulated. The accumulation of toxic concentrations of H_2_S is prevented by its oxidation to thiosulfate and sulfate via the mitochondrial sulfide oxidation pathway (12). The first and committed step in this pathway is catalyzed by sulfide quinone oxidoreductase (SQR), an inner mitochondrial membrane-anchored flavoprotein, which is a member of the flavin disulfide reductase superfamily (13). SQR couples H_2_S oxidation to coenzyme Q_10_ (CoQ_10_) reduction (12,14–16). It transfers the oxidized sulfane sulfur to a small molecule acceptor, which is predicted to be glutathione (GSH) under physiological conditions (15,16). Inherited deficiency of SQR presents as Leigh disease (17).

While the overall reaction catalyzed by SQRs are similar (18–20), the requirement of a small molecule acceptor by the human enzyme distinguishes it from bacterial homologs, which build long polysulfide chains and can release octasulfur rings as oxidation products (18–20). In contrast, the catalytic cycle of human SQR resembles that of bacterial flavocytochrome *c* sulfide dehydrogenase, which couples the conversion of sulfide to hydrodisulfide with the reduction of cytochrome *c* (21). Unexpectedly, the crystal structures of human SQR revealed the presence an additional sulfur bridging the active site cysteines in a trisulfide (22,23). The catalytic relevance of the trisulfide configuration is controversial, and it has been assigned as the inactive (22) or active (23) form of the enzyme. The presence of a cysteine trisulfide in SQR raises questions about how it is built in the active site. To our knowledge, a catalytically relevant cysteine trisulfide would be the first of its kind for a thiol-based redox active cofactor.

A reaction mechanism for human SQR that starts with the trisulfide as the resting form of the enzyme is shown in Fig. 1. The reaction cycle proceeds via two half reactions. In the first half reaction, sulfide adds to the trisulfide at the solvent-accessible Cys-379 to form a ^379^Cys-SSH persulfide. The bridging sulfur is retained on ^201^Cys-SS^-^ persulfide, which forms an unusually intense charge transfer (CT) complex with FAD that is centered at 695 nm (14,16,23,24). Sulfur transfer from ^379^Cys-SSH to small molecule acceptor leads to regeneration of the active site trisulfide with the concomitant two-electron reduction of FAD. In the second half reaction, FADH2 transfers electrons to CoQ_10_, regenerating the resting enzyme and linking sulfide oxidation to mitochondrial energy metabolism by supplying reduced CoQ_10_ to Complex III in the electron transport chain (25).

**Figure 1.**
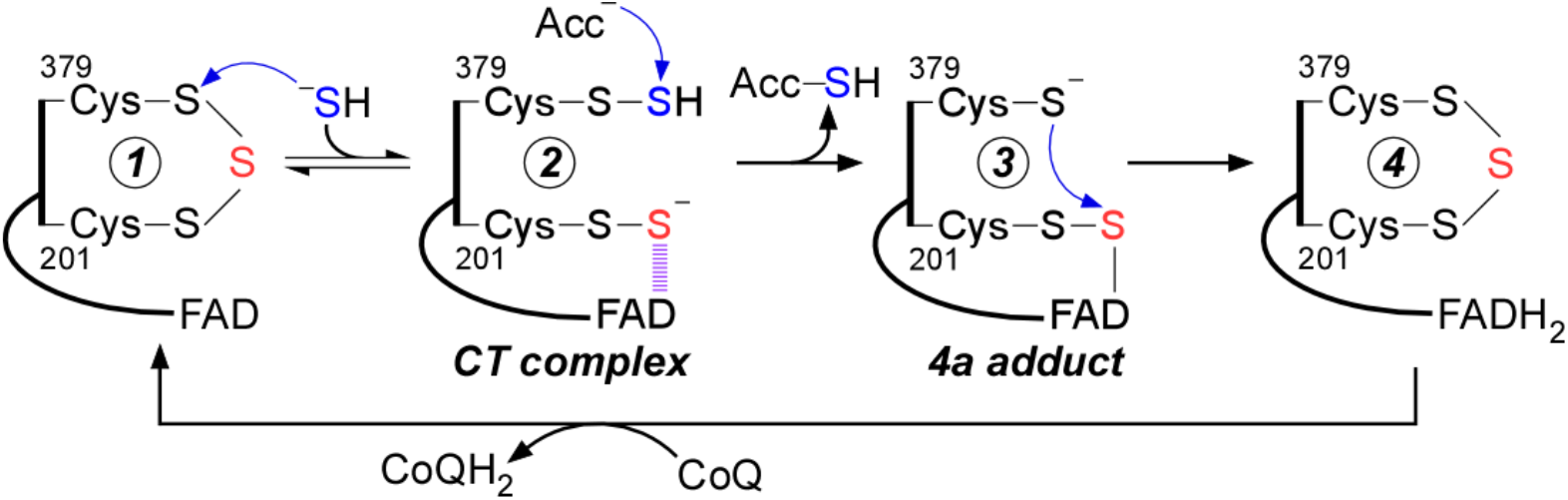
Postulated mechanism for sulfide oxidation catalyzed by human SQR. Proposed mechanism for the reaction catalyzed by SQR. Sulfide adds into the resting cysteine trisulfide (1) to generate a ^379^Cys-SSH persulfide and a ^201^Cys-SS^-^ persulfide, with the latter participating in a CT complex with FAD (2). Sulfur transfer to a small molecule acceptor proceeds through a putative 4a adduct (3) to generate the reduced enzyme (4). Electron transfer from FADH2 to CoQ regenerates the resting enzyme. The oxidized sulfur and bridging sulfur in the cysteine trisulfide are labeled in blue and red, respectively.

In principle, an active site cysteine trisulfide provides several advantages over the conventional disulfide configuration seen in the mechanistically similar flavocytochrome c sulfide dehydrogenase (21). Sulfane sulfur species have increased electrophilic character versus their respective thiols (26), which would enhance the reactivity of the solvent-accessible Sγ of Cys-379 in SQR towards nucleophilic addition by sulfide. Indeed, the rate of sulfide addition to the cysteine trisulfide of SQR is estimated to be ~2Í10^7^- fold higher than the rate of sulfide addition to cysteine disulfide in solution (0.6 M^-1^s^-1^ at pH 7.4, 25 °C) (27). The subsequent formation of persulfide rather than thiolate intermediate on Cys-201 would also enhance its reactivity for facilitating sulfur transfer and electron movement via the putative C4a adduct.

In this study, we report the spectral and kinetic characterization of cyanolysis-induced dismantling followed by sulfide-dependent rebuilding of the trisulfide cofactor. Cyanide treatment destabilized human SQR and led to its inactivation with concomitant loss of the bridging sulfane sulfur. Addition of sulfide to inactive cyanide treated enzyme led to recovery of active SQR, indicating that the oxidation state of the active site cysteines was preserved upon cyanide treatment. Crystallization of SQR with cyanide led to the capture of a ^379^Cys *N*-(^201^Cys-disulfanyl)-methanimido thioate intermediate, providing insights into how the trisulfide can be rebuilt following cyanide treatment. Finally, computational modeling indicated that the trisulfide configuration provides a significant catalytic advantage over a disulfide in the SQR reaction. Collectively, our study demonstrates that the cysteine trisulfide in SQR is required for its catalytic activity, confers a catalytic edge over a disulfide, and contributes to its structural integrity.

## Results

### Formation and decay of the cyanide-induced CT complex in SQR

Mixing SQR with cyanide led to the formation of an intense CT complex characterized by an absorbance maximum at 695 nm and a shift in the FAD peak from 450 nm to 420 nm, with isosbestic points at 430 nm and 505 nm (Fig. 2A). These spectral features are similar to the CT complexes seen previously with other nucleophiles (24,28). From the dependence of the rate of CT complex formation on the concentration of cyanide, the following parameters were obtained: *k*_on_ = 10,500 ± 118 M^-1^ s^-1^, *k*_off_ = 3.7 ± 0.6 s^-1^, and *K*_D(app)_ = 348 ± 52 μM at 4 °C (Fig. 2B,C). The CT complex is an intermediate in the catalytic cycle of SQR, and sulfide addition to the CT intermediates formed by alternative nucleophiles leads to their decay with the concomitant reduction of FAD (24,28). Similarly, addition of sulfide immediately following cyanide-induced CT complex formation led to its decay with concurrent reduction of FAD (Fig. 2D). This result indicates that the CT complex formed in the presence of cyanide can participate in the first half reaction leading to FADH2 formation.

**Figure 2.**
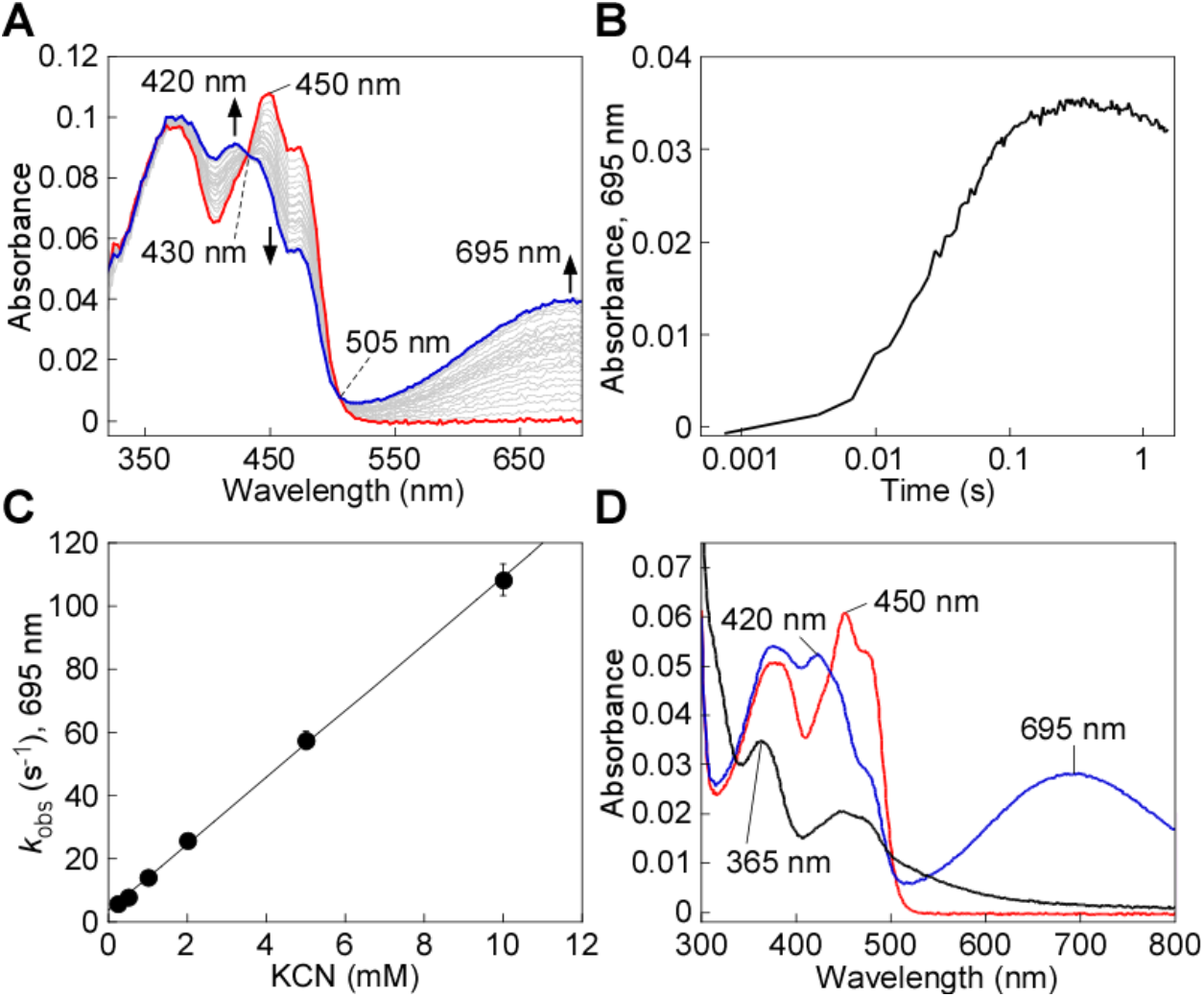
Cyanide-induced CT complex formation in SQR. **A**, SQR (10 μM, red line) in 100 mM potassium phosphate, pH 7.4 containing 0.03% DHPC, was mixed 1:1 (v/v) with KCN (4 mM) and monitored over 1.5 s at 4 °C for the formation of the cyanide-induced CT complex at 695 nm (blue line). **B**, Representative stopped flow kinetic trace for the reaction in (A) monitored at 695 nm. **C**, Dependence of the *k*_obs_ at 4 °C for cyanide-induced CT complex formation on cyanide concentration. The data are representative of two independent experiments, with each data point obtained in triplicate. **D**, SQR (5 μM, red line) was treated with KCN (5 mM) to form the CT complex (blue line), immediately followed by the addition of Na2S (200 μM) and incubated for 5 min at 20 °C, which led to CT complex decay and FAD reduction (black line). The data are representative of three independent experiments.

### Decay of the cyanide-induced CT complex and cyanolysis of the cysteine trisulfide

Extended incubation of the cyanide-induced CT complex in the presence of excess cyanide led to its slow decay (Fig. 3A). A *k*_obs_ of 0.15 ± 0.02 min^-1^ at 20 °C was observed for the decay of the CT complex in the presence of 5-10 mM KCN (Fig. 3B). The FAD spectrum following CT decay was slightly altered from that in native SQR. Thus, a blue shift in the absorbance maximum from 450 nm to 447 nm and a narrowing of the 380 nm absorption peak were seen (Fig. 3A). The altered spectral features were observed even after the enzyme was desalted to remove excess cyanide, suggesting a change in the flavin electronic environment. Cyanolysis yields ~1 mol of sulfane sulfur per mol SQR monomer (23), indicating that the native trisulfide state is lost upon prolonged cyanide treatment. To confirm this conclusion, we added sulfite, which forms a strong CT complex when added to native SQR (23), However, sulfite did not elicit spectral changes in cyanide pre-treated and desalted SQR (Fig. 3C).

**Figure 3.**
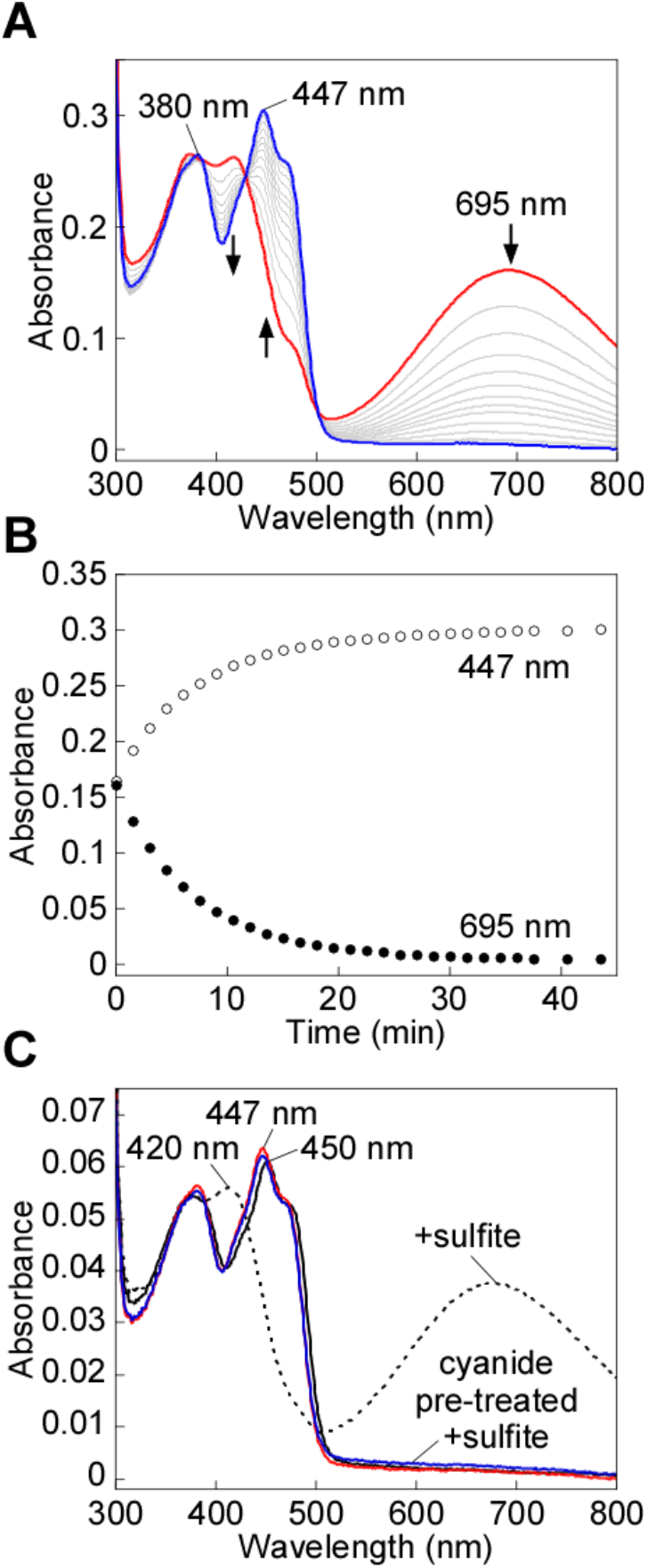
Cyanide-induced CT complex decay in SQR. **A**, SQR (25 μM) in Buffer A was treated with KCN (10 mM) to form the CT complex (red line), which was monitored over 43 min at 20 °C for the complete decay of the CT complex (blue line). **B**, Kinetic traces for the decay of the cyanide-induced CT complex in (A), monitored at 450 nm (open circles) and 695 nm (closed circles). **C**, SQR (5 μM, solid black line) was treated with sodium sulfite (5 mM) and incubated for 1 min at 20 °C to form the CT complex formation (dashed black line). In tandem, SQR (5 μM) pre-treated with KCN (10 mM) and desalted (solid red line) was then treated with sodium sulfite (5 mM). CT complex formation was not observed after incubation for 1 min at 20 °C (solid blue line). The data are representative of three independent experiments.

### Sulfide-mediated regeneration of the active site trisulfide

We next assessed the impact of cyanide treatment on SQR activity under steady state turnover conditions. Surprisingly, the specific activity of SQR in the standard assay was similar for the cyanide pre-treated (369 ± 25 μmol min^-1^ mg^-1^) and native (360 ± 12 μmol min^-1^ mg^-1^) enzymes. This result suggested that cyanide treated enzyme could be reactivated by rebuilding the trisulfide following cyanolysis of SQR.

We therefore monitored the rate at which the trisulfide is rebuilt using as a measure of the active enzyme, formation of the sulfide-induced CT complex (14,24) (Fig. 4A). For this, the kinetics of CT complex formation was assessed following rapid mixing of sulfide with cyanide pre-treated SQR. Compared to native SQR (*k*_obs_ = 19.4 ± 1.8 s^-1^), CT complex formation was ~12-fold slower with cyanide pre-treated SQR (1.6 ± 0.2 s^-1^) (Fig. 4B). The lag in the absorbance increase at 675 nm indicated that trisulfide rebuilding limits the rate of CT complex formation in cyanide pre-treated SQR. Consistent with this postulate, incubation of cyanide pre-treated SQR with sulfide for 1 h at 4 °C led to FAD reduction (Fig. 4C), signaling reformation of the active trisulfide-containing SQR under these conditions. The presence of excess sulfide, which serves as both the sulfur donor and acceptor, led to FADH2 accumulation in the absence of CoQ_1_ (14,24). The 447 nm FAD absorption peak observed in cyanide pre-treated SQR (Fig. 2A) shifted to 450 nm following incubation with sulfide (Fig. 4D), indicating recovery of the native FAD microenvironment. Furthermore, addition of sulfite to regenerated SQR resulted in the formation of a robust CT complex, confirming the presence of a trisulfide in the active site (Fig. 4D).

**Figure 4.**
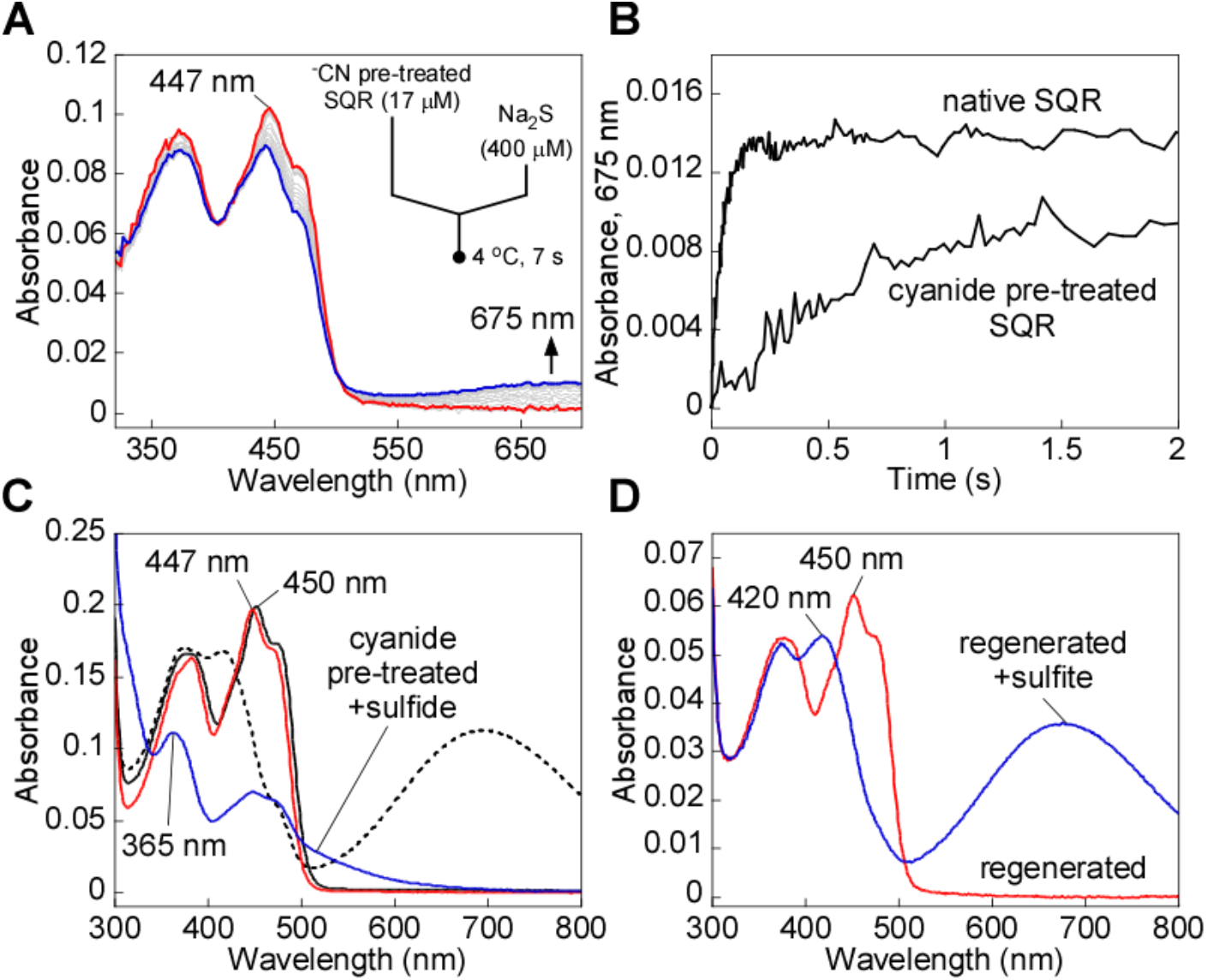
Regeneration of cyanide pre-treated SQR by sulfide. **A**, Cyanide pre-treated SQR (17 μM, red line) in Buffer A was rapidly mixed 1:1 (v/v) with Na2S (400 μM) and monitored over a period of 7 s at 4 °C for formation of the sulfide-induced CT complex (blue line). **B**, Comparison of the kinetic traces at 675 nm for cyanide pre-treated SQR, as shown in A, versus native SQR mixed with Na2S (400 μM) under the same conditions. **C**, SQR (17 μM, solid black line) in Buffer A was treated with KCN (10 mM) to form the CT complex (dashed black line) and monitored over 40 min at 20 °C for the complete decay of the CT complex and desalted to remove excess cyanide (red line). Cyanide pre-treated SQR was then incubated with Na2S (300 μM) for 1 h at 4 °C, which led to FAD reduction (blue trace). **D**, Cyanide pre-treated SQR, pre-incubated with sulfide under the same conditions as (A) and desalted (5 μM, red line), was treated with sulfite (5 mM) and incubated for 1 min to form the sulfite-induced CT complex (blue line). The data are representative of three independent experiments.

### Cyanolysis of the bridging sulfur decreases SQR protein stability

We consistently observed that cyanide treatment led to an increased tendency for SQR to aggregate at temperatures above 20 °C, indicating that loss of the bridging sulfur in the active site trisulfide leads to protein instability. We therefore investigated the thermal stability of SQR with and without cyanide pre-treatment. Native SQR exhibited a Tagg of 64.8 °C, compared to 36.5 °C for cyanide pre-treated enzyme (Fig. 5). Incubation of cyanide pre-treated SQR in the presence of excess sulfide increased its stability (Tagg of 56.6 °C). Thus, the decrease in thermal stability of SQR upon loss of the bridging sulfur was largely reversed upon regeneration of the cysteine trisulfide.

**Figure 5.**
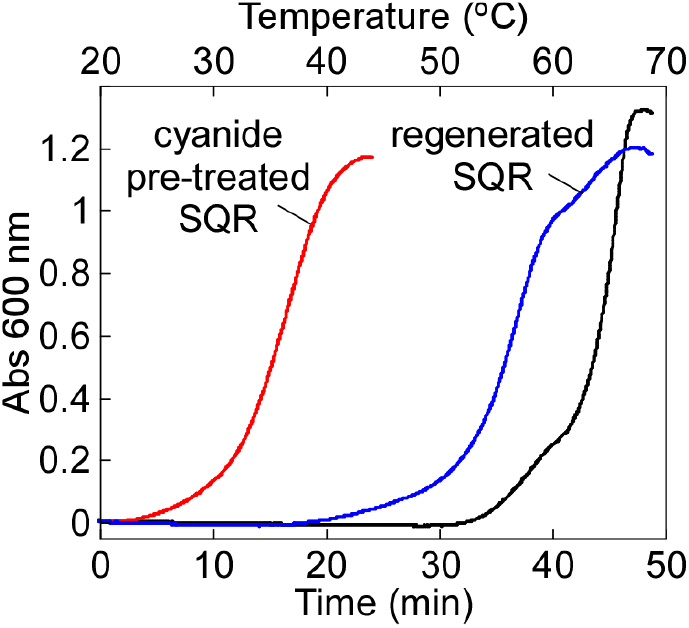
Effect of bridging sulfur extraction on SQR protein stability. SQR (20 μM) in Buffer A was pre-treated with KCN (10 mM) for 45 min at 20 °C and desalted, followed by incubation with Na2S (300 μM) for 1 h at 4 °C and a second desalting. A final SQR concentration of 5 μM was used for the thermal denaturation assays. The stability of native SQR (black line) versus cyanide pre-treated SQR before (red line) and after (blue line) incubation with sulfide was monitored by the increase in absorbance at 600 nm. The data are representative of three independent experiments.

### Dithiol-mediated reduction of FAD in SQR

As an alternative to cyanolysis, we attempted to extract the bridging sulfur from the SQR trisulfide using DTT, which in principle could reduce the trisulfide to generate free thiols on Cys-379 and Cys-201. Unexpectedly, treatment with DTT led to bleaching of the yellow color associated with SQR (Fig. 6A). Given the known substrate promiscuity of SQR (24,28), we postulate that DTT adds to the resting trisulfide, forming a mixed disulfide and a CT complex. In the second step, an intramolecular displacement by the second thiol in the ^379^Cys-S-S-DTT adduct leads to elimination of oxidized DTT, reduction of FAD, and regeneration of the trisulfide (Fig. 6B).

**Figure 6.**
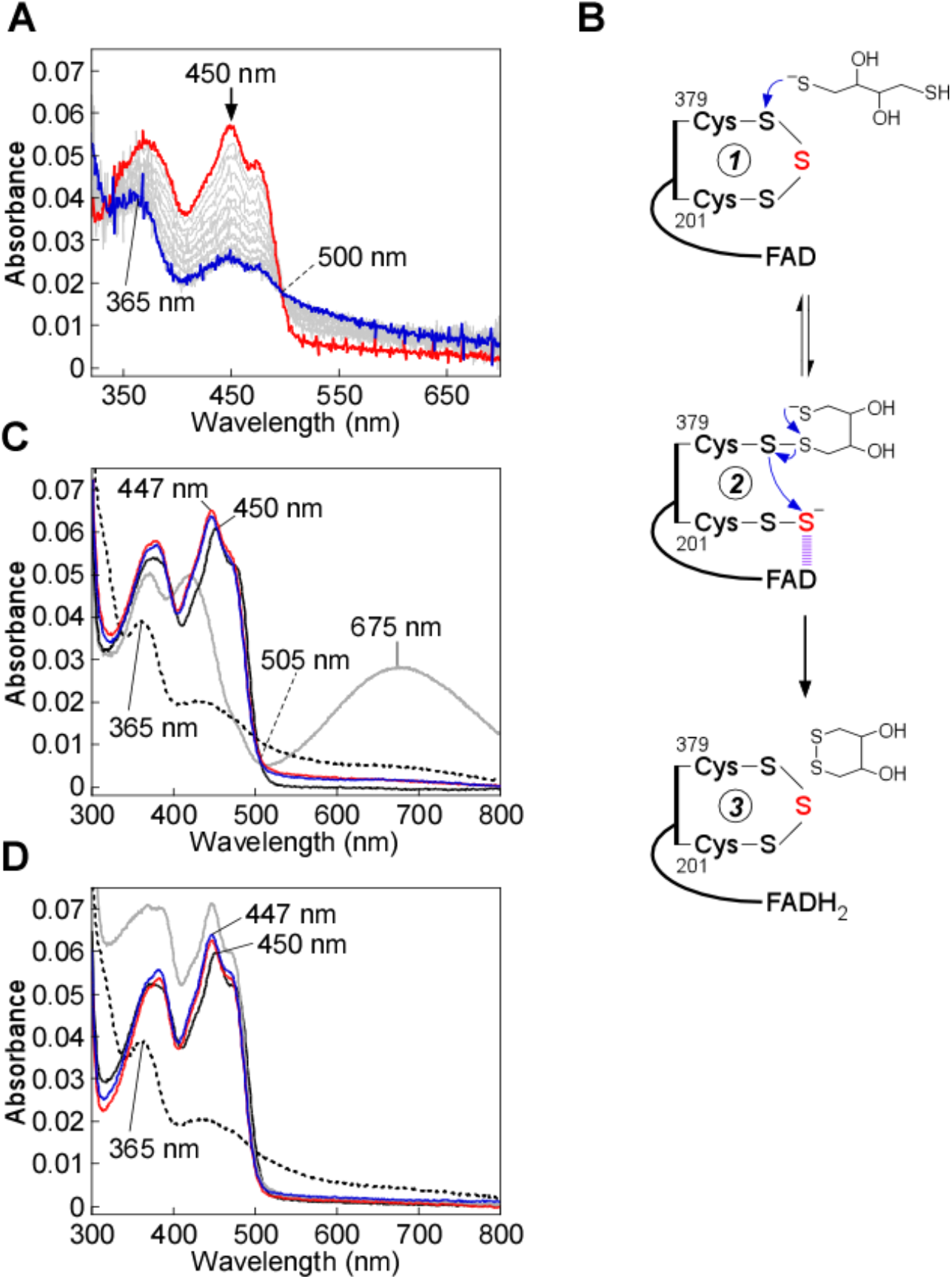
Dithiol-mediated reduction of FAD in SQR. **A**, SQR (10 μM) in Buffer A was rapidly mixed 1:1 (v/v) with DTT (400 μM) and FAD reduction was monitored over 7 s at 4 °C. **B**, Proposed mechanism for the addition of DTT into the SQR cysteine trisulfide, leading to FAD reduction. DTT adds into the cysteine trisulfide at the solvent-accessible Cys-379 to generate a mixed disulfide and ^201^Cys-SS^-^. An intramolecular thiol-disulfide exchange then regenerates the SQR cysteine trisulfide, with electrons moving into FAD. **C**, SQR (5 μM) in Buffer A (solid black line), was treated with DTT (200 μM), leading to FAD reduction (dashed black line), or β-mercaptoethanol (200 μM), leading to stable CT complex formation (solid gray line). FAD reduction was not observed in cyanide pre-treated SQR (5 μM, solid red line) upon treatment with DTT (200 μM, solid blue line). **D**, SQR (5 μM) under the same conditions as in (C) (solid black line), was treated with DHLA (200 μM), leading to FAD reduction (dashed black line), followed by re-oxidation by addition of CoQ_1_ (180 μM, solid gray line). FAD reduction was not observed in cyanide pre-treated SQR (5 μM, solid red line) upon treatment with DHLA (200 μM, solid blue line). The data are representative of three independent experiments.

To further test this model, the reaction of DTT with SQR was characterized by stopped-flow spectroscopy. Upon mixing SQR rapidly with DTT, FADH2 formation (*k*_obs_ = 0.36 ± 0.04 s^-1^) was observed without accumulation of a CT complex intermediate (Fig. 6A). This contrasted with the reaction of other nucleophiles with SQR, and suggested that resolution of the mixed disulfide, via an intramolecular reaction, is more rapid than its formation (Fig. 6B). Next, we tested whether the intact cysteine trisulfide in SQR is required for FAD reduction by DTT. Pre-treatment of SQR with cyanide prevented FAD reduction by DTT (Fig. 6C).

In contrast to DTT, the monothiol, β-mercaptoethanol, was unable to drive FAD reduction, and formed a stable CT complex instead (Fig. 6C). Like DTT, the dithiol dihydrolipoic acid (DHLA), a physiological reductant, also led to FAD reduction, but only when the cysteine trisulfide was intact (Fig. 6D).

### Structure of SQR-CoQ_1_ soaked with cyanide

To obtain structural insights into the interaction of cyanide with SQR, crystals of human SQR-CoQ_1_ were soaked with cyanide. The 2.25 Å resolution structure was obtained by molecular replacement using coordinates for the SQR-CoQ_1_ structure (PDB ID: 6OIB) (Table 1). The overall structure (Fig. 7A) is similar to that reported previously for SQR-CoQ_1_ (23).

**Figure 7.**
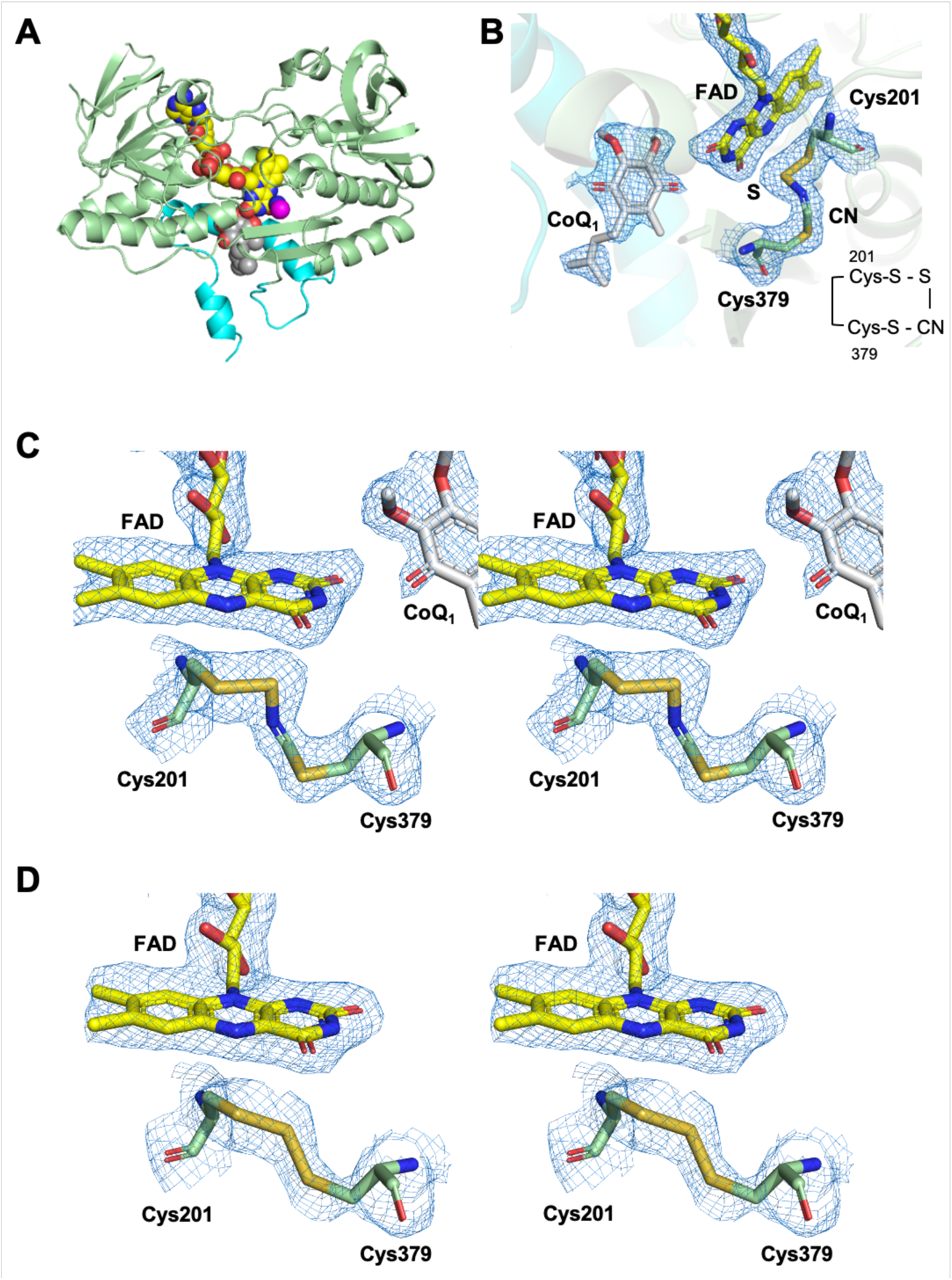
Structure and active site of SQR-CoQ_1_ + cyanide. **A**, The overall structure of SQR-CoQ_1_ + cyanide is shown with FAD, CoQ_1_, and cyanide in yellow, grey, and magenta spheres, respectively. The C-terminal membrane-anchoring helices are highlighted in cyan. **B**, Electron density maps (2F_o_-F_c_) of the active site shown in mesh contoured at 1.0 σ. Cys-201, Cys-379, FAD, CoQ_1_, sulfur derived from the trisulfide, and cyanide are shown in stick display. **C**, Stereo image of the active site of SQR-CoQ_1_ treated with cyanide. The electron densities (2F_o_-F_c_) are contoured at 1.0 σ**. D**, Stereo image of the active site in SQR-CoQ_1_ treated with sulfide (PDB ID: 6OI6) showing the resting trisulfide. Chain A is shown in this figure.

**Table 1.** Crystallographic data collection and refinement statistics^*^

|  | **SQR-CoQ<sub>1</sub> with cyanide** |
| --- | --- |
| Space group | P2<sub>1</sub>2<sub>1</sub>2<sub>1</sub> |
| Unit cell parameters (Å) | a=78.39<br>b=111.75<br>c=134.05<br>$\alpha=\beta=\gamma=90^\circ$ |
| Wavelength (Å) | 1.12723 |
| **<u>Data collection statistics</u>** |  |
| Resolution range (Å) | 50.00-2.25 (2.29-2.25) |
| Number of unique reflections | 57050 (2696) |
| Completeness (%) | 99.7 (96.0) |
| R<sub>merge</sub> | 0.175 (0.974) |
| R<sub>pim</sub> | 0.059 (0.359) |
| Redundancy | 8.6 (6.6) |
| Mean I/σ | 14.0 (2.3) |
| **<u>Refinement statistics</u>** |  |
| Resolution range (Å) | 39.20-2.24 |
| R<sub>work</sub>/R<sub>free</sub> (%) | 17.24/21.60 |
| RMSD bonds (Å) | 0.008 |
| RMSD angles (deg) | 0.922 |
| Average B factor (Å<sup>2</sup>) | 36.62 |
| Number of water molecules | 125 |
| Ramachandran |  |
| favored (%) | 96.51 |
| allowed (%) | 3.49 |
| not allowed (%) | 0 |

\*Values in parentheses are for highest-resolution shell.

Surprisingly, strong and continuous electron density was observed between Cys-201 and Cys-379 (Fig. 7B,C). The electron density in the presence of cyanide was more extended than for the trisulfide in the native SQR-CoQ_1_ + sulfide structure (Fig. 7D). We interpret the additional electron density as evidence for the insertion of a cyanide molecule in the trisulfide bridge, forming an *N*-(^201^Cys-disulfanyl)-methanimido thioate intermediate, ^201^Cys-S-S-N=CH-S-^379^Cys. The relevance of this species to the spectral intermediates observed in the presence of cyanide is discussed later.

### MD simulations and QM/MM reactivity predictors for a disulfide versus trisulfide cofactor

Direct comparison of the active site architecture in representative SQR structures was extracted from 600 ns trajectories (RMSDs shown in Fig. S1). The simulations provided insights into the structural and dynamical differences between the trisulfide versus a modeled disulfide state, and a chemical rationale for the use of the trisulfide cofactor by SQR. In the disulfide structure (Fig. 8A, r*ight*) the sulfur atoms of Cys-201 and Cys-379 are buried (Fig. S2, *right*), and not in contact with solvent molecules. The Sg atoms in the two cysteines exhibit differences in their distance to the C4a in FAD, with Cys-379 being closer (Fig. 8B, *right*) at distances of 3.4 Ǻ versus 3.9 Ǻ for Cys-201 in the representative structure. These data argue against a catalytic disulfide configuration in SQR, which is reinforced by considerations of the intrinsic electrophilicity of the sulfur atoms calculated at the QM/MM level.

**Figure 8.**
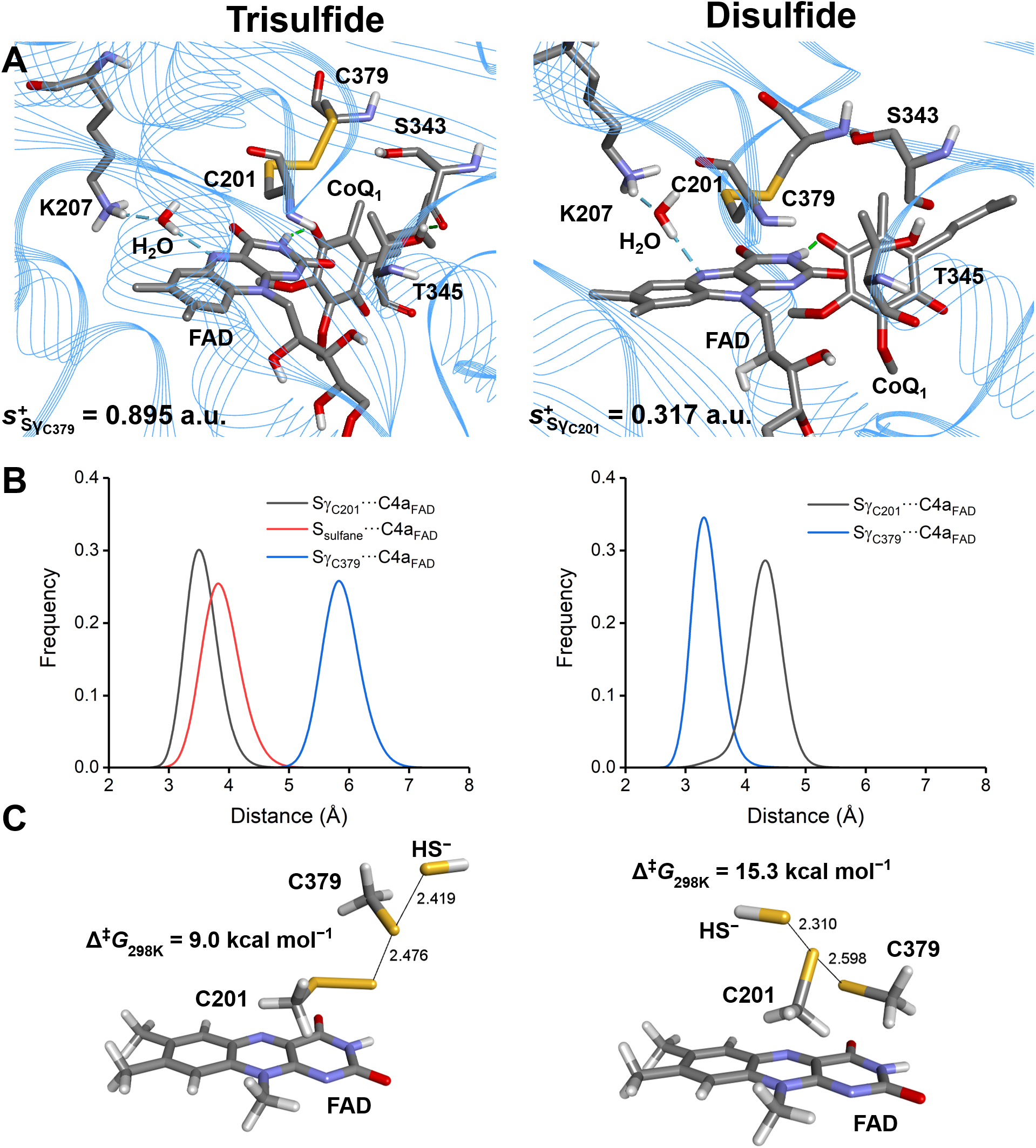
MD simulations and computational modeling of SQR. **A,** Active site architecture in representative structures corresponding to the most populated cluster from 600 ns MD simulations of SQR in the trisulfide (left) or disulfide (right) state. Condensed local softness for the most electrophilic Sγ atom between Cys-201 and Cys-379 is reported for each system in atomic units (a.u.). **B,** Sulfur-to-C4a FAD distances for S_γC201_–C4a_FAD_/S_sulfane_-C4a_FAD_/S_γC379_–C4a_FAD_ (trisulfide, left) and S_γC201_-C4a_FAD_/S_γC379_–C4a_FAD_ (disulfide, right) monitored along the corresponding MD trajectories. **C,** Structure of the transition states (TS) located for the sulfide anion attack on the trisulfide (left) or disulfide (right) using a reduced model of the active site of SQR at the IEFPCM-DFT level of theory in a dielectric of *ε* = 10.125. Data correspond to interatomic distances in Å and Gibbs free-energy associated barriers at 298 K in kcal mol^−1^.

In the trisulfide model, the Sg atom of Cys-201 is estimated to be intrinsically less electrophilic than Cys-379 (s^+^ = 0.317 versus 0.895 atomic units). These results support the proposed attack of the sulfide anion on the Sg atom of Cys-379 as the first step in the catalytic mechanism (Fig. 1). In the trisulfide structure (Fig. 8A, *left*) the sulfur atom of Cys-379 is located in a small cavity and is solvent exposed (Fig. S2, *left).* The sulfane sulfur in the trisulfide points inward, sits at the apex of a 109° S-S-S angle, and is almost equidistant from the C4a atom in FAD as the Sg of Cys-201. The 4.9 Å (sulfane sulfur) and 4.2 Å (Sg of Cys-201) distances to C4a in FAD in the representative structure from the most populated cluster in solution (Fig. 8A, *left*), are comparable to the 4.3 and 3.3 Å distances observed in the trisulfide-containing crystal structure of SQR-CoQ_1_ + sulfide (22). Inspection of the distribution of values for SC201-C4aFAD and for the sulfane sulfur-C4aFAD distances along the simulation (Fig. 8B, *left*) reveal broader histograms compared to the disulfide ones and a 0.5 Å difference between the maxima.

The atomic charges calculated for the sulfur atoms in Cys-201, Cys-379 and the sulfane sulfur in the trisulfide, reveal a slightly electropositive reaction zone, particularly over Cys-379 (+0.112 atomic units Table S1), favoring attack of the negatively charged sulfide anion. On the other hand, the local softness for gaining electron density reveals that the Sg of Cys-379 and the sulfane sulfur are similar and significantly more reactive than the Sg of Cys-201.

### Density Functional Theory (DFT) in a Polarized Continuum Model (PCM) characterization of the sulfide addition step

To gain further insights into the specific reactivities of the electrophilic sulfurs in a disulfide (Cys-201) versus a trisulfide (Cys-379) cofactor, we modeled the detailed mechanism using a DFT/PCM level of theory as described under Experimental Procedures. The SN2 mechanism involved the attack of a sulfide anion on the trisulfide or a hypothetical disulfide, with a persulfide or a thiolate anion, respectively serving as the leaving group (Fig. 8C). The transition states are quasi-linear, late, and quite synchronic, both in terms of heavy atom reorganization (HAR) and CT complex, with HAR/CT complex being more advanced in the disulfide compared to the trisulfide, as evidenced by the Wiberg Bond indices (WBI) and natural population analysis (NPA) charges (Tables S2-S5). The computed free energy barriers for the reaction of sulfide anion with the trisulfide versus disulfide cofactor in SQR are 9.0 and 15.3 kcal mol^-1^ at 25 °C and 1 atm, respectively. These results provide strong supporting evidence for the significantly greater reactivity of the trisulfide over the disulfide, accounting for much of the 10^7^-fold difference in the second order rate constant for the reaction of sulfide anion with SQR versus with a disulfide in solution [23]. The structure of the resulting CT product complex (Fig. S4) confirms completion of the SN2 reaction, and that the persulfide (or thiolate) can proceed to the next step in the reaction mechanism, concentrating excess negative charge at the ^201^Cys-SS^-^ or (^379^Cys-S^-^).

## Discussion

Members of the flavoprotein disulfide reductase superfamily have a signature two redox cofactor active site constellation. While the flavin is common to all members, the second cofactor can be a cysteine disulfide, which is the most common theme in the superfamily, a cysteine sulfenic acid, or a mixed disulfide (e.g. Cys-S-S-CoA) (13). Recently, a fourth variation on the redox active cysteine cofactor theme, i.e., a trisulfide, was discovered in human SQR (22,23).

Examples of cysteine trisulfides in proteins are rare. They have been observed primarily as artifacts in recombinant human growth hormone preparations (29–32). Trisulfide intermediates have been postulated as catalytic intermediates in dissimilatory sulfite reduction in bacteria (33) and in the SQR-catalyzed polysulfide formation in *Aquifex aeolicus* (18). The role of the cysteine trisulfide in human SQR is controversial (22,23). Initially, it was proposed to result from a dead-end reaction with sulfide under anaerobic conditions in the absence of a sulfur acceptor (14). In this model, the active site cysteine disulfide in SQR would be regenerated via a chemically unusual mechanism that necessitates the elimination of Sγ of Cys-379 as an oxidized product, and replaces it with the sulfur atom derived from the trisulfide bridge (22). In addition to the unusual chemistry, the mechanism would require a significant conformational change to shorten the ~3.5 Å distance between the Sγ of Cys-379 and Cys-201 to allow formation of a cysteine disulfide. Recent biochemical data from our laboratory have however, indicated that the trisulfide in SQR likely represents the active form of the enzyme (23). In this study, we disassembled and then reassembled the active site trisulfide by cyanolysis followed by sulfuration, and demonstrated that these processes led to the restoration of SQR activity.

The presence of the cysteine trisulfide in SQR was previously confirmed biochemically via cold cyanolysis (23), which extracts the bridging sulfur as thiocyanate (34). During the cyanolysis reaction, we had observed a rapid color change from yellow to blue, followed by a slow reversion to yellow, indicating the transient formation of a cyanide-induced CT complex followed by its decay. In the current study, this mechanism was supported by spectral and kinetic analyses, which demonstrate that cyanide, acting as a nucleophile, adds into the cysteine trisulfide (Fig. 2). We propose that cyanide attacks at the solvent-accessible Cys-379, forming a ^379^Cys-S-C≡N organic thiocyanate and a ^201^Cys-SS^-^ persulfide-to-FAD CT complex (Fig. 9, *2*). The intense CT complex is similar to those seen with alternative nucleophiles such as sulfite or methanethiol adding to human SQR (24,28) and also resembles the CT complex induced by coenzyme A persulfide in short-chain acyl-CoA dehydrogenase (23). The off-rate constant (*k*_off_ = 3.7 ± 0.6 s^-1^ at 4 °C) indicates that cyanide-induced CT complex formation is reversible, and that cyanide dissociation regenerates the trisulfide. Notably, cyanide can also act as a sulfur acceptor in the SQR reaction forming thiocyanate (14), and can thus contribute to the sulfide-mediated FAD reduction when both sulfide and cyanide are present. The ability of cyanide treated SQR to support catalysis indicated that the oxidation status of the active site cysteines was preserved in cyanolyzed enzyme.

**Figure 9.**
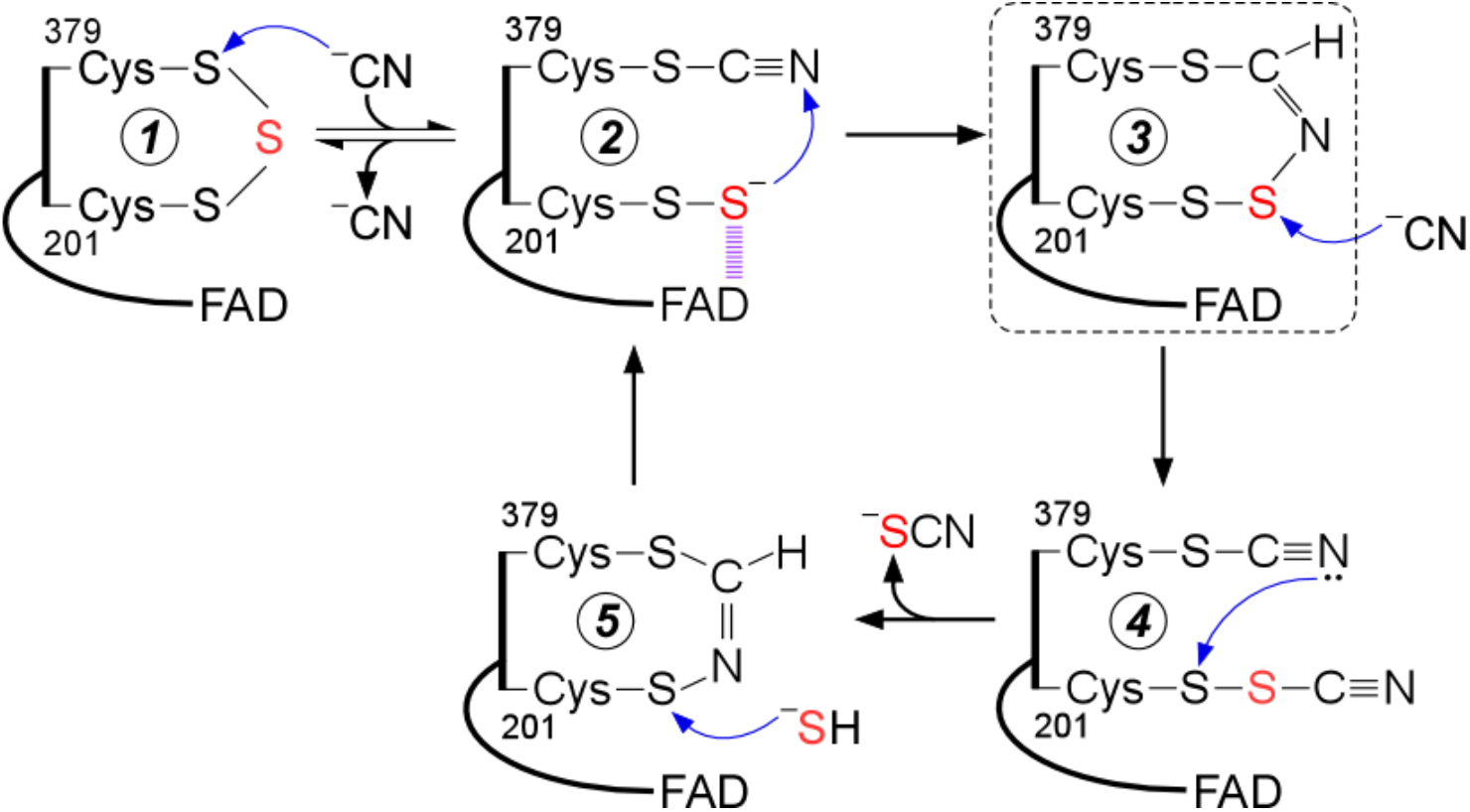
Proposed mechanism for cyanolysis and cysteine trisulfide rebuilding in SQR. Cyanide adds into the resting cysteine trisulfide (1) to generate a ^379^Cys-S-CN organic thiocyanate while the bridging sulfur is retained in the ^201^Cys-SS^-^ persulfide that participates in a CT complex with FAD (2). Conversion to the ^379^Cys *N*-(^201^Cys-disulfanyl)-methanimido thioate intermediate (3) leads to loss of the CT complex. Addition by a second cyanide at the sulfane sulfur of Cys-201 leads to intermediate (4), which can cyclize and eliminate thiocyanate (5), completing the cyanolysis reaction. Addition of sulfide to the Sγ of Cys-201 in the ^379^Cys *N*-(^201^Cys-sulfanyl)-methanimido thioate intermediate (5) regenerates the CT complex (2). Elimination of cyanide regenerates the resting trisulfide form of the enzyme. The bridging sulfur of the cysteine trisulfide is labeled in red. The dashed box highlights the intermediate observed in the crystal structure.

The crystal structure of SQR provided a clue as to how the redox state of the cysteines is maintained upon cyanide treatment by revealing a bridging ^379^Cys *N*-(^201^Cys-disulfanyl)-methanimido thioate intermediate (Fig. 7). We propose that this intermediate is formed by attack of the ^201^Cys-SS^-^ persulfide on the ^379^Cys-S-C≡N thiocyanate (Fig. 9, *3*). While the bridging *N*-(disulfanyl)-methanimido thioate intermediate is stabilized *in crystallo*, it is susceptible to attack by a second equivalent of cyanide (Fig. 9, *4*), leading to thiocyanate elimination, which was detected by the cold cyanolysis reaction. The resulting ^201^Cys-S-N=CH-S-^379^Cys intermediate (Fig. 9, *5*) preserves the redox state of the active site cysteines. It does not however, support generation of a sulfite-induced CT complex (Fig. 3C), or FAD reduction in the presence of dithiols (Fig. 6 C, D), and it destabilizes SQR (Fig. 5). The absorption spectrum of FAD in the presence of this intermediate is subtly different from that in the native enzyme with the 450 nm peak blue shifted to 447 nm, and the 380 nm peak being better resolved.

We propose that the trisulfide is rebuilt by the nucleophilic attack of sulfide on the Sγ of Cys-201, leading to a CT complex and then, to the resting enzyme (Fig. 9, 5→2→1). We attribute the lag phase that was seen by stopped flow spectroscopy when sulfide was mixed with cyanide treated versus untreated SQR, to the time needed to rebuild the active enzyme trisulfide (Fig. 4A, B). Once rebuilt, the enzyme cycles through multiple catalytic turnovers and a difference in specific activities was not seen in cyanide treated versus untreated SQR under steady-state assay conditions.

The difference in the active site configurations of untreated versus cyanide treated SQR was further demonstrated by their differential reactivity to the dithiols DTT and DHLA. Both dithiols can substitute for sulfide in the oxidative half reaction, transferring electrons to FAD to form FADH2 (Fig. 6). Neither DTT nor DHLA reduced FAD in cyanide pretreated SQR, supporting the proposed mechanism (Fig. 6B).

Computational QM/MM and QM modeling provide strong evidence for the catalytic relevance of the trisulfide versus the disulfide form of the cofactor in SQR (Fig. 8). Based on accessibility, electrostatics and local softness considerations, the combination of MD simulations and QM/MM modeling predicted that Cys-379 in the trisulfide is the electrophilic target in the first step of SQR mechanism. Based on DFT/PCM modeling of the first step in the reaction mechanism, i.e. the attack by a sulfide anion, it was estimated that the trisulfide configuration affords an ~10^5^-fold rate enhancement over a disulfide cofactor in the active site of SQR.

A significant question raised by the discovery of the trisulfide in SQR is how is the cofactor built? Minimally, one of two mechanisms can be considered. In the first, both cysteines are oxidized (e.g. to a sulfenic acid) followed by the attack of a sulfide anion to form a persulfide (e.g. on the solvent accessible Cys-379), setting up trisulfide formation. In the second mechanism, both cysteines are persulfidated, an oxidative cysteine modification that has been detected in many proteins (35). Low molecular weight persulfides (e.g. cysteine persulfide) could lead to formation of the bis-persulfide form of SQR from which the trisulfide could be built. Low molecular weight persulfides can be synthesized by all three H_2_S generating enzymes (6,7,36,37), and a potential role for these reactive sulfur species in signaling has been suggested (38). Alternatively, generation of the trisulfide could be catalyzed; candidate human sulfur transferases include rhodanese (39), mercaptopyruvate sulfurtransferase (6) and TSTD1 (40).

In summary, we have demonstrated that the cysteine trisulfide in human SQR can be reversibly dismantled and reassembled. The trisulfide not only contributes to a significant rate enhancement over a disulfide for the nucleophilic addition of sulfide, but also stabilizes the enzyme. Studies are underway in our laboratory to investigate whether assembly of the trisulfide is enzyme catalyzed.

## Materials and Methods

### Materials

The following reagents were purchased from Millipore Sigma: CoQ_1_, n-dodecyl-β-D-maltoside (DDM), potassium cyanide, sodium sulfide nonahydrate, and sodium sulfite. The phospholipids, DHPC (1,2-diheptanoyl-*sn*-glycero-3-phosphocholine) and POPC, were purchased from Avanti Polar Lipids (Alabaster, AL). DHLA was purchased from Cayman Chemical Company (Ann Arbor, MI).

### Preparation of human SQR

Human SQR was purified as detergent-solubilized recombinant enzyme as described previously (15). Human SQR used for crystallization was purified in an identical procedure as described previously (15), except that DHPC (0.03% w/v) was substituted with DDM (0.05% w/v) as the solubilizing detergent (23).

### SQR spectral analyses and activity assays

The absorption spectra of SQR were recorded on a temperature-controlled Shimadzu UV-2600 spectrophotometer in Buffer A (50 mM Tris, pH 8.0, containing 300 mM NaCl and 0.03% DHPC). The concentration of SQR used in the spectral assays was estimated by the absorbance of the FAD cofactor, using an extinction coefficient of 11,500 M^-1^ cm^-1^ at 450 nm (14). SQR activity was estimated by the rate of CoQ_1_ reduction (Δ_εox-red_ = 12,000 M^-1^cm^-1^) at 25 °C as described previously (15), using sulfite (800 μM) as the sulfur acceptor.

### Stopped flow spectroscopy

All stopped flow experiments were conducted at 4 °C on a SF-DX2 double mixing stopped-flow system from Hi-Tech Scientific, equipped with a photodiode array detector (300-700 nm range). The concentrations reported in the figure legends for stopped flow experiments are before 1:1 (v/v) mixing.

### Detection of sulfane sulfur in SQR

SQR was assayed for sulfane sulfur using the cold cyanolysis method as described previously (23). The data for mol sulfane sulfur per mol SQR monomer are presented as the mean ± SD of three independent preparations of SQR.

### Crystallization of SQR-CoQ_1_ with cyanide

SQR-CoQ_1_ crystals were grown at 20 °C by the hanging drop vapor diffusion method using solubilized human SQR (17.4 mg mL^-1^) in 50 mM Tris-HCl pH 8.0, containing NaCl (300 mM) and n-dodecyl β-D-maltoside (0.05% w/v) supplemented with CoQ_1_ (5 mM, in 100% DMSO). The SQR solution was then mixed 1:1 (v/v) with the reservoir solution composed of 200 mM ammonium tartrate dibasic, pH 6.6, and PEG 3350 (20% w/v), yielding a final CoQ_1_ concentration of 2.5 mM. The resulting SQR-CoQ_1_ crystals were soaked with potassium cyanide (1.25 mM) for 40 min, followed by cryoprotection in the aforementioned reservoir solution supplemented with glycerol (35% v/v) before freezing in liquid nitrogen.

### Thermal denaturation assays

The thermal stabilities of untreated SQR, and cyanide pre-treated SQR before and after sulfide treatment, were assessed using 300 μL of enzyme (5 μM) in a quartz cuvette, housed in a temperature-controlled Shimadzu UV-2600 spectrophotometer. SQR was allowed to equilibrate at 20 °C for 2 min before initiating the assay by increasing the temperature by 1 °C min^-1^. Thermal denaturation was monitored by the increase in absorbance at 600 nm.

### X-ray data collection and structure determination

Diffraction data for SQR-CoQ_1_ + cyanide crystal was collected at the LS-CAT beamline 21-ID-D (Advanced Photon Source, Argonne National Laboratory) at 1.12723 Å wavelength. The diffraction images were processed using HKL2000 (41). The molecular replacement solution for SQR-CoQ_1_ + cyanide was determined using SQR-CoQ_1_ (PDB ID: 6OIB) as a search model. The final structures were completed using alternate cycles of manual fitting in Coot (42) and refinement in REFMAC5 (43). The stereochemical quality of the final models was assessed using MolProbity (44).

### MD simulations of SQR

The crystal structure of human SQR-CoQ_1_ + sulfide in the trisulfide state complexed with FAD (PDB: 6OI6, monomer A, 2.56 Ǻ resolution) was used as a starting point (23). CoQ_1_ was manually docked by superimposing another structure of SQR-CoQ_1_ (PDB: 6OIB, monomer A, 2.03 Å resolution). Two systems were simulated: SQR in a Cys-201-Cys-379 disulfide state (SQR-SS) and in the trisulfide state (SQR-SSS). Protonation and tautomers of titratable residues, and missing hydrogen atoms, were added with the ProToss utility (45). Systems were solvated with a periodic truncated octahedral box of TIP3P water extended up to 12 Å around solute, then neutralized with six Cl^-^ ions with the *leap* utility of AmberTools17 (46). Both were then minimized, heated to 310 K (500 ps, NVT), and equilibrated at 1 atm (1 ns, NPT) prior to conducting the simulations (600 ns, NPT). Minimization and simulation was carried out using the *pmemd.cuda* module of AMBER 16 (46). For describing the protein, the AMBER *ff14SB* force field was used for standard residues, whereas the *gaff* force field was used for FAD and CoQ_1_ ligands, along with RESP charges (47). The central sulfur atom of the trisulfide moiety was treated as a separate residue with zero charge. All parameters for describing the trisulfide moiety were present in the *ff14SB* force field, except for the S-S-S bonds, which were taken from the *gaff* force field. An 8.0 Å cutoff was used for treating direct non-bonding interactions, and long-range interactions were treated with the *Particle Mesh Ewald* (PME) method (48). For MD simulations, temperature and pressure (in NPT simulations) were controlled by means of the Langevin thermostat (49) and the Monte Carlo barostat (50), respectively. Distances involving hydrogen atoms were constrained with SHAKE (51) and a time integration step of 2 femtoseconds was used. Harmonic restraints of 10 kcal mol^-1^ Å^-2^ were applied to a water-bridged hydrogen bond observed in SQR crystal structures (23) between the protonated amino group of the Lys-207 side chain and the N5 atom in FAD. Trajectory processing and analysis was done with the *cpptraj* module of AmberTools17 (46). Convergence of simulations was monitored following the Cα-RMSD (see Fig. S1). In order to extract representative structures from the MD 600 ns trajectories, clustering analysis (5 clusters) was performed for each system, using a hierarchical-agglomerative algorithm with *cpptraj* (46).

### Reactivity descriptors from QM/MM calculations on SQR

Descriptors of intrinsic reactivity derived from the electronic structure of Cys-379 and Cys-201 were calculated at the QM/MM level in the framework of a conceptual DFT (52) at the M06-2X-D3/6-31+G(d,p) level of theory (53–55) combined to a classical description of the enzyme using the aforementioned force fields. Global softness (*S)* and electrophilic Fukui function condensed to the Sg atom 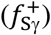 were calculated according to equations (1) and (2):

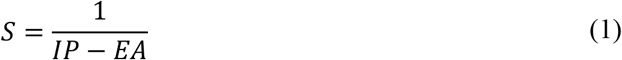

where *IP* and E*A* respectively represent the ionization potential and electron affinity of the system of interest, determined using the vertical DSCF (self-consistent field) approximation (56), and

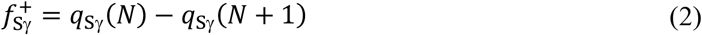

where *q*_Sg_(*N)* and *q*_Sg_(*N*+1) represent the atomic charge on the S_γ_ atoms of Cys-379/Cys-201, calculated using a Natural Population Analysis (57), both in the system of reference bearing N electrons and after addition of one extra electron. The atomic electrophilic softness (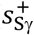, calculated as *S* times 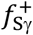) is a local descriptor that can be used to compare Sg intrinsic reactivity in Cys-379 and Cys-201 across SQR-SS and SQR-SSS. The electronic structure of each macromolecular system was thus obtained through *single-point* calculations performed on representative structures extracted from MD simulations using the additive QM/MM scheme implemented in AMBER16 (46) interfaced with Gaussian 09 Rev. D.01 (58) with a QM region comprising ^379^Cys-CH2-S(S)S-CH2-^201^Cys.

### DFT-PCM modeling of reaction mechanisms and barriers for sulfide nucleophilic attack

The mechanism of the reaction of the sulfide anion, manually docked and oriented as guided by our previous models of similar reactions (59), was characterized at the M06-2X-D3/6-31+G(d,p)-PCM level of theory, previously validated by us to model reactions of the sulfide anion in a similar system (59). A simplified representation of the catalytic disulfide/trisulfide and FAD at the active site of SQR was used including CH_3_SSSCH_3_/ CH_3_SSCH_3_ and the flavin. The structures of each reactant complex, transition and product complex were fully optimized and verified by the inspection of the eigenvalues of the Hessian matrix at the same level. Thermochemical corrections at 298 K and 1 atm were calculated under usual approximations of statistical thermodynamics (rigid rotor, harmonic frequencies) as implemented in Gaussian 09 Rev. D01 (58). The effects exerted by the bulk protein on the active site structure along the reaction and reaction barrier were introduced using the IEF-PCM continuum model (60) with a dielectric constant ε = 10.125. The reactive systems were placed in a molecular shaped cavity constructed using Bondi’s radii (61) and including non-electrostatic (cavitation, repulsion and dispersion) contributions. In order to connect transition states with reactant complexes and product complexes, we calculated the IRC reaction path (62) using the HPC algorithm (63).

## Acknowledgements

We thank Drs. David P. Ballou and Markus Ruetz for insightful discussions on the cysteine trisulfide cyanolysis and rebuilding mechanisms. MD simulations were carried out using Uruguayan supercomputer resources from ClusterUY (https://cluster.uy). Continuous support from PEDEClBA-Uruguay is gratefully acknowledged by JB and ELC who are active members of the Uruguayan National Research System (SNI-ANII).

## Author contributions

A.P.L. designed and performed the kinetic and spectroscopic experiments. S.M. determined the crystal structure, which was analyzed together with U.S.C. J.B. performed the computational modeling and MD simulations, which were analyzed together with E.L.C. R.B. helped conceive the experiments, analyzed the data and co-wrote the manuscript with A.P.L. and S.M. with the exception of the computational sections that were co-written by J.B. and E.L.C. All authors approved the final version of the manuscript.

## Abbreviations used

H_2_S: hydrogen sulfide
CoQ_10_ or CoQ_1_: coenzyme Q_10_ or Q_1_
CT: charge transfer
SQR: sulfide quinone oxidoreductase
GSH: glutathione
DTT: dithiothreitol
DHLA: dihydrolipoic acid
HAR: heavy atom reorganization
MD: molecular dynamics
QM/MM: hybrid quantum mechanics/molecular mechanics electronic structure modeling
DFT: density functional theory
PCM: polarized continuum model.

