## Supplemental Figures S1-S3, Tables S1-S5 for "Dismantling and rebuilding the trisulfide cofactor demonstrates its essential role in human sulfide quinone oxidoreductase"

### Supporting Figures and Tables

Aaron P. Landry[1], Sojin Moon[1], Jenner Bonanata[2], Uhn Soo Cho[1], E. Laura Coitiño[2] and Ruma Banerjee[1]*

[1]Department of Biological Chemistry, University of Michigan Medical School, Ann Arbor, MI 48109 and [2]Laboratorio de Química Teórica y Computacional (LQTC), Instituto de Química Biológica, Facultad de Ciencias and Centro de Investigaciones Biomédicas (CeInBio), Universidad de la República, Iguá 4225, Montevideo 11400, Uruguay

**Table of contents**

Figure S1-S3
Tables S1-S5

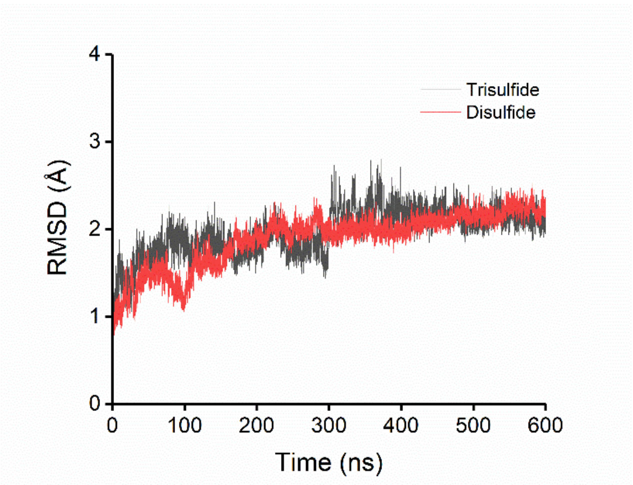


**Figure S1.** Temporal evolution of Cα-RMSD along the 600 Amber MD simulations of SQR in the trisulfide (dark gray) and the disulfide (red) states, confirming convergence.

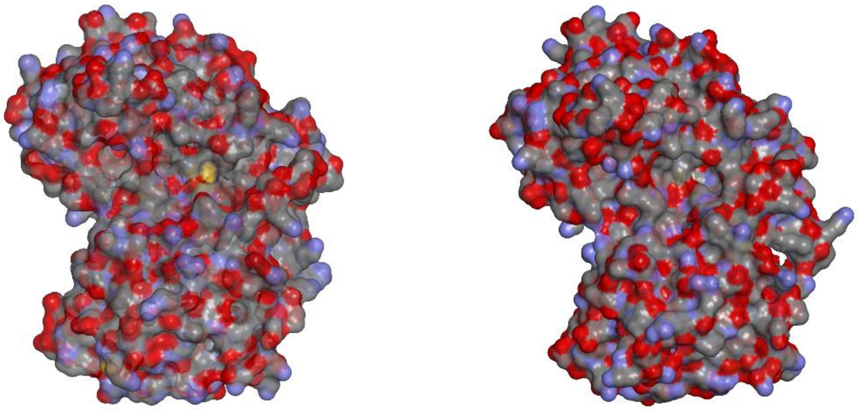

**Figure S2.** The solvent accessibility of Cys-201/Cys-379 is compared in representative structures of the most populated cluster from MD simulation for trisulfide (left, most populated cluster, c0, 39%) versus disulfide (right, most populated cluster, c0, 41%) states. The molecular surface of SQR is colored by atom type (C, gray; N, blue; O, red; S, yellow).

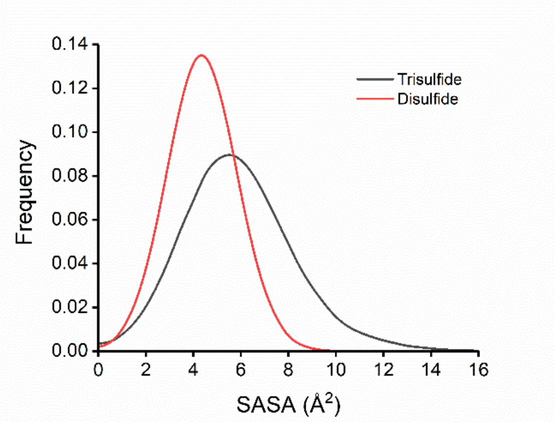


**Figure S3.** Histograms for the solvent accessible surface area (SASA in Å$^2$) of the more exposed sulfur in SQR trisulfide (Sγ Cys-379, black) and in SQR-disulfide (Sγ C201, red) models long the 600 nsec MD simulations.

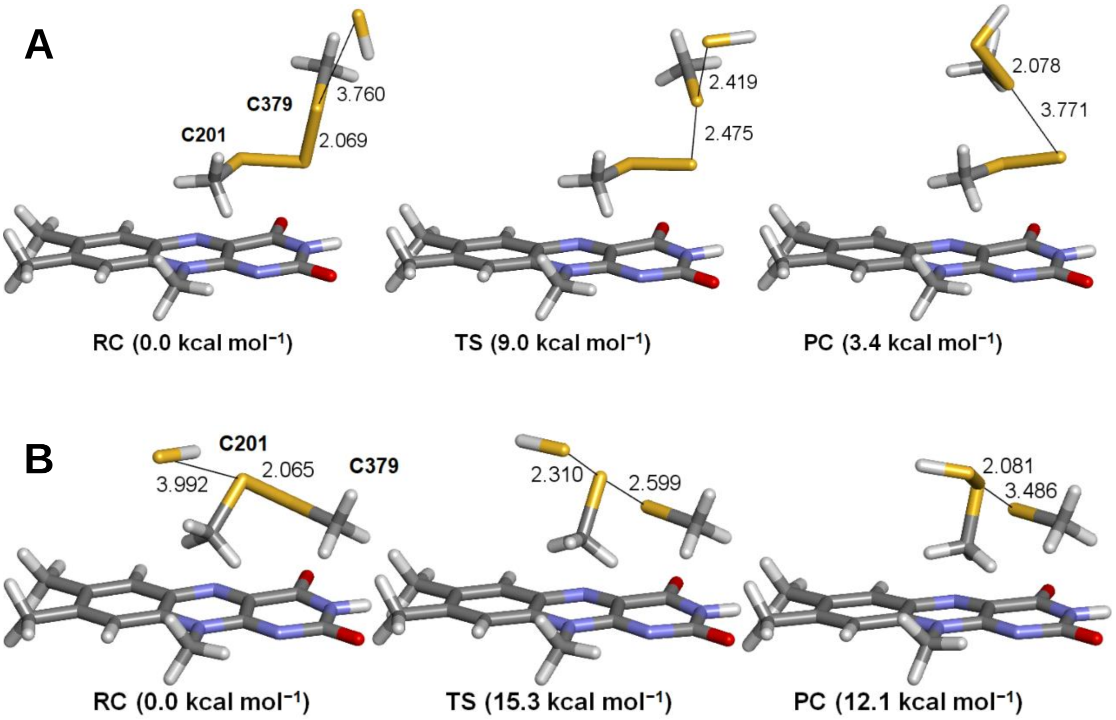


**Figure S4.** Structure of the species located at the PCM-M06-2X-D3/6-31+G(d,p) level of theory in a dielectric of ε = 10.125 for the attack of the sulfide anion on trisulfide (A) and disulfide (B) using a reduced model of the active site of SQR. RC, TS and PC represent the reactant complex, transition state and product complex, respectively. Data corresponds to interatomic distances in Å and relative Gibbs free-energy at 298 K in kcal mol$^{-1}$ calculated with respect to the RC.

**Table S1.** Predictors of intrinsic reactivity calculated at the M06-2X-D3/6-31+G(d,p)/AMBER level on complete SQR in the trisulfide form, in atomic units.

| Atom (k) | $q_k^{+a}$ | $f_k^{+b}$ | $s_k^{+c}$ |
|---|---|---|---|
| S$\gamma_{C201}$ | +0.032 | 0.150 | 0.446 |
| $S_{sulfane}$ | −0.011 | 0.318 | 0.946 |
| S$\gamma_{C379}$ | +0.112 | 0.301 | 0.895 |

[a] Natural population analysis (NPA) atomic charges on atom k; [b] condensed Fukui electrophilic function for atom k; [c] local softness for electrophilic reactivity at atom k.

**Table S2.** Wilberg bond indices (WBI, a.u.) for relevant pairs of atoms and their evolution along the species participating in the attack of a sulfide anion on the trisulfide modeled at the IEFPCM-M06-2X-D3/6-31+G(d,p) level of theory in a dielectric of ε = 10.125.

| Atom pair | RC | TS | PC |
|---|---|---|---|
| $S_{SH}$–S$\gamma_{C379}$ | 0.013 | 0.559 | 1.024 |
| S$\gamma_{C379}$–$S_{trisulfide}$ | 1.020 | 0.475 | 0.015 |
| $S_{trisulfide}$–S$\gamma_{C201}$ | 1.014 | 1.031 | 1.033 |
| $S_{trisulfide}$–C4$X_{FAD}$ | 0.002 | 0.004 | 0.023 |
| $S_{trisulfide}$–C$9_{FAD}$ | 0.002 | 0.003 | 0.009 |
| S$\gamma_{C201}$–C4$X_{FAD}$ | 0.008 | 0.013 | 0.008 |
| S$\gamma_{C201}$–C$9_{FAD}$ | 0.006 | 0.008 | 0.006 |

**Table S3.** NPA atomic charges (in atomic units) for relevant atoms and their evolution along the species participating in the attack of a sulfide anion on the trisulfide in SQR modeled at the IEFPCM-M06-2X-D3/6-31+G(d,p) level of theory in a dielectric of ε = 10.125.

| Atom | RC | TS | PC |
|---|---|---|---|
| $S_{SH}$ | −1.11 | −0.61 | −0.22 |
| S$\gamma_{C379}$ | +0.09 | +0.03 | +0.07 |
| $S_{trisulfide}$ | −0.06 | −0.42 | −0.78 |
| S$\gamma_{C201}$ | +0.07 | +0.02 | −0.01 |
| C4$X_{FAD}$ | +0.09 | +0.10 | +0.10 |
| N$5_{FAD}$ | −0.34 | −0.35 | −0.36 |

Sum of NPA charges on flavin group in PC: –0.09 atomic units.

**Table S4.** WBI (atomic units) for relevant pairs of atoms and their evolution along the species participating in the attack of the sulfide anion on the disulfide modeled at the IEFPCM-M06-2X-D3/6-31+G(d,p) level of theory in a dielectric of ε = 10.125.

| Atom pair | RC | TS | PC |
|---|---|---|---|
| $S_{SH}$–S$\gamma_{C201}$ | 0.007 | 0.679 | 1.019 |
| S$\gamma_{C201}$–S$\gamma_{C379}$ | 1.036 | 0.370 | 0.016 |
| S$\gamma_{C379}$–C4$X_{FAD}$ | 0.007 | 0.020 | 0.036 |
| S$\gamma_{C379}$–C$9_{FAD}$ | 0.007 | 0.012 | 0.016 |

**Table S5.** NPA atomic charges (in atomic units) for relevant atoms and their evolution along the species participating in the attack of the sulfide anion on the disulfide using modeled at the IEFPCM-M06-2X-D3/6-31+G(d,p) level of theory in a dielectric of ε = 10.125.

| Atom | RC | TS | PC |
|---|---|---|---|
| $S_{SH}$ | −1.10 | −0.50 | −0.22 |
| $S\gamma_{C201}$ | +0.06 | +0.04 | +0.08 |
| $S\gamma_{C379}$ | +0.05 | −0.45 | −0.71 |
| $C4X_{FAD}$ | +0.08 | +0.11 | +0.11 |
| $N5_{FAD}$ | −0.33 | −0.36 | −0.37 |

Sum of NPA charges on flavin group in PC: –0.11 atomic units.
